# Reconstitution of basic mitotic spindles in cell-like confinement

**DOI:** 10.1101/770602

**Authors:** Sophie Roth, Ioana C. Gârlea, Mathijs Vleugel, Bela M. Mulder, Marileen Dogterom

## Abstract

Bipolar organization of the mitotic spindle is the result of forces generated by dynamic microtubules and associated proteins in interaction with chromosomes and the cell boundary^1–4^. Biophysical experiments on isolated spindle components have provided important insights into the force-generating properties of different components^5–8^, but a quantitative understanding of the force balance that results from their concerted action is lacking. Here we present an experimental platform based on water-in-oil emulsion droplets that allows for the bottom-up reconstitution of basic spindles. We find a typical metaphase organization, where two microtubule asters position symmetrically at moderate distance from the mid-zone, is readily obtained even in the absence of chromosomes. Consistent with simulations, we observe an intrinsic repulsive force between two asters that can be counterbalanced alternatively by cortical pulling forces, anti-parallel microtubule crosslinking, or adjustment of microtubule dynamics, emphasizing the robustness of the system. Adding motor proteins that slide anti-parallel microtubules apart drives the asters to maximum separation, as observed in cells during anaphase^9,10^. Our platform offers a valuable complementary approach to *in vivo* experiments where essential mitotic components are typically removed, instead of added, one by one.

---

The mitotic spindle consists of a dynamic assembly of microtubules (MTs), motor proteins, and other microtubule associated proteins (MAPs), which serves to segregate duplicated chromosomes during mitosis. The molecular parts list of a typical mammalian spindle includes structural proteins, force-generating components as well as a large number of proteins that regulate the spindle through the subsequent phases of mitosis (**Fig. 1a**)^1–4^. A large body of work in tissue culture cells has provided us with important insights into the roles of different spindle components, and biophysical experiments on isolated components have taught us about the magnitude of forces that can be generated.

**Figure 1.**
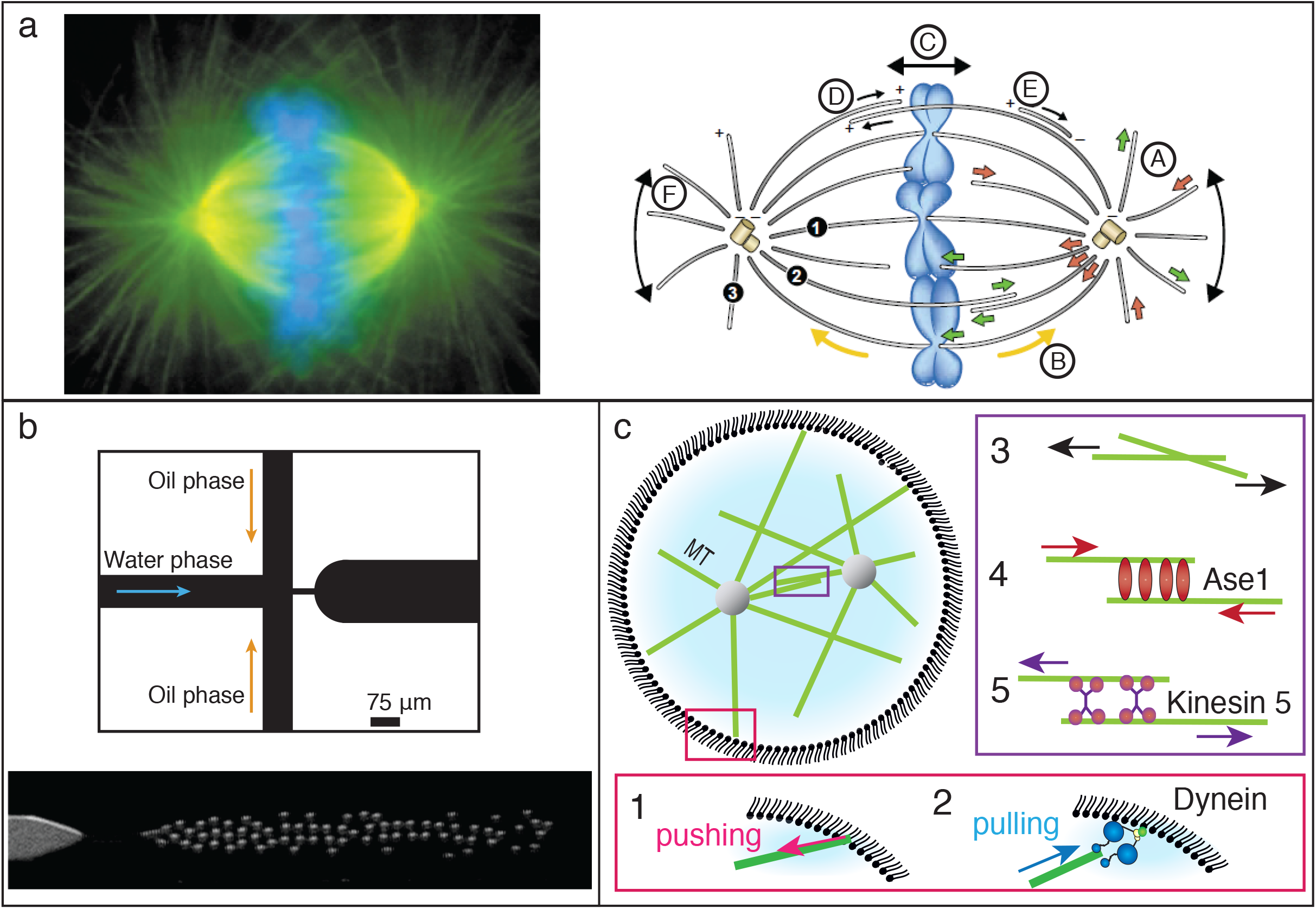
Forces in mitotic spindles. **(a)** Left: Static immunofluorescence photograph of a mitotic spindle in a tissue-culture cell. Green, MTs; blue, chromosomes. Right: The spindle contains different subpopulations of MTs: kinetochore MTs (1), interpolar MTs (2) and astral MTs (3). A variety of dynamic processes occur in the spindle, including MTs undergoing dynamic instability (A, green and red arrows represent growing and shrinking MTs); poleward MT flux (B, yellow arrows); chromosome movements (C); motor-driven antiparallel MT sliding (D); dynein dependent, minus-end directed MT transport (E); and orientational movements of the spindle poles (F) (reproduced with permission from Wittmann et al.^46^). **(b)** Flow focusing microfluidic chip, as described in^32, 33^, used to generate W/O emulsion droplets with a diameter in the range 12-25μm. Top: Flow focusing junction. Bottom: Fluorescent image of dextran FITC in the water phase. **(c)** Schematic representation of the force generating spindle components studied in this paper. Two dynamic asters are confined in a W/O emulsion droplet. At the boundary, growing MTs generate pushing forces^47^ (1) and MTs interacting with cortical dynein exert pulling forces^5^ (2). In the overlap zone, interdigitating MTs generate steric repulsive forces^48^ (3), Ase1 molecules which crosslink antiparallel MTs exert entropy-driven attractive forces between the centrosomes^30^ (4), Kinesin 5 molecular motors which slide antiparallel MTs apart, generate repulsive forces between the centrosomes^6^ (5).

Spindle positioning and orientation result from the complex interplay between forces generated within the spindle and at the cortex, and are directly influenced by the shape of the cell^11–13^. Dynamic MTs growing against rigid structures such as the cell cortex can exert length-dependent pushing forces^14–16^. On the other hand, when MT ends are captured by the molecular motor dynein at the cell cortex, shrinking MTs can generate pN pulling forces^5^. Assembly of a bipolar spindle is also directly affected by motor proteins that slide interpolar, anti-parallel MTs, generating inward or outward forces^17–21^. In particular, MT plus-end directed motors of the kinesin-5 family generate outward forces that drive bipolarity^6, 9, 22–29^. In contrast, diffusible crosslinkers from the MAP65 family (PRC1 in vertebrate and Ase1 in yeast) generate friction that prevents MT separation^7^. The entropic forces generated by these crosslinkers are sufficient to antagonize motor protein-driven MT sliding *in vitro*^30^, and are thought to play a crucial role in the overall force balance within the spindle.

The contributions of these components to the overall force balance and resulting spindle organization have typically been inferred from top-down perturbations of *in vivo* spindles in which components of interest are individually depleted. However, interpretation of the outcome of such experiments is inherently difficult given the complexity of the cellular context. Here, we reconstitute minimal spindles starting from two centrosome-nucleated microtubule asters within the cell-like confinement of water in oil (W/O) emulsion droplets (**Fig. 1b**). We use this system to directly probe the contributions of dynamic MTs (pushing forces and steric interactions), dynein-mediated cortical pulling forces, anti-parallel crosslinkers, and anti-parallel motor proteins to spindle organization (**Fig. 1c**).

In order to mimic the geometrical confinement in which spindles assemble and orient in a vertebrate mitotic cell, we generated spherical water in oil emulsion droplets of cell-like size (15-25μm), lined by a single lipid monolayer. Within these droplets, we first asked how spindle positioning is affected by pushing and pulling forces generated by MTs at the cortex. We studied both the assembly and positioning of single confined dynamic asters (**Fig. 2**), and the behaviour of two such asters inside droplets (**Fig. 3**). Dynamic MTs were categorised into three regimes, based on whether the average MT length was shorter (1), approximately equal (2) or longer (3) than the radius *R* of the droplet corresponding to early, intermediate and late stages in our experiments.

**Figure 2.**
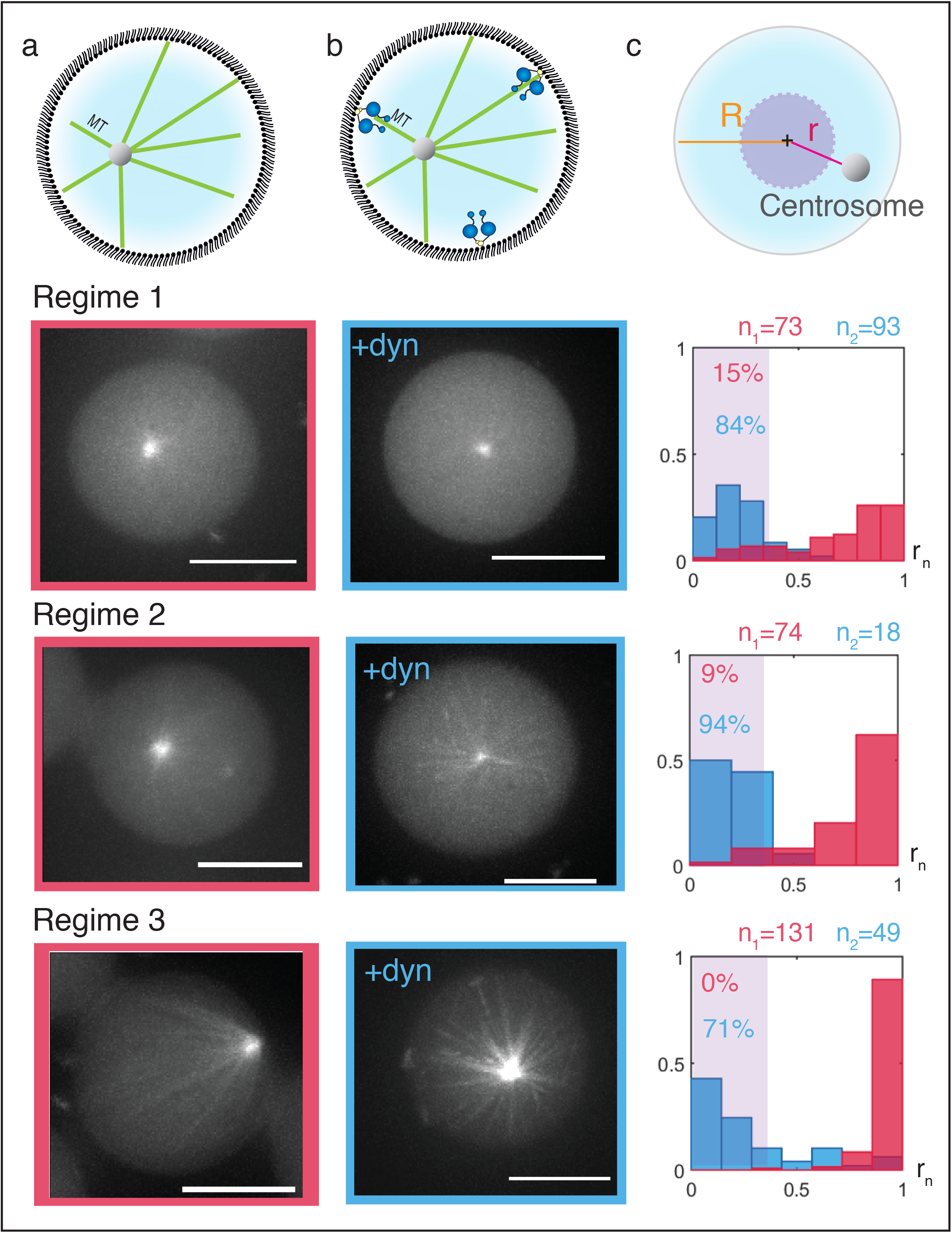
Single aster positioning by pushing and pulling forces at the cortex. **(a)(b)** Positioning of one dynamic aster in absence (a) or presence (b) of dynein at the cortex. Top: Schematic picture. Bottom: Spinning disk confocal fluorescence images for regimes 1, 2 and 3 where the average MT length is respectively short, intermediate and long compared to the radius of the droplet (Tubulin is fluorescently labelled, Scale=10μm). **(c)** Quantification of aster position. Top: Representation of r, the distance from centrosome to centre and R, the radius of the droplet. The purple sphere is concentric with the droplet and represents 5% of the droplet volume. Bottom: Histograms of centrosome position 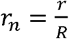 in the absence (pink) or presence (blue) of dynein at the R cortex. The bin size was defined as 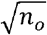 with n_0_ = min (*n*_*1*_, *n*_*2*_), and *n*_*1*_, *n*_*2*_ is the total number of asters quantified in absence or presence of dynein respectively. The number of asters N found in each bin is normalized. The purple zone indicates centred asters. The percentage of centred asters in the absence (pink) or presence (blue) of dynein is indicated. In the absence of dynein, asters adopt a peripheral position as MTs grow.

**Figure 3.**
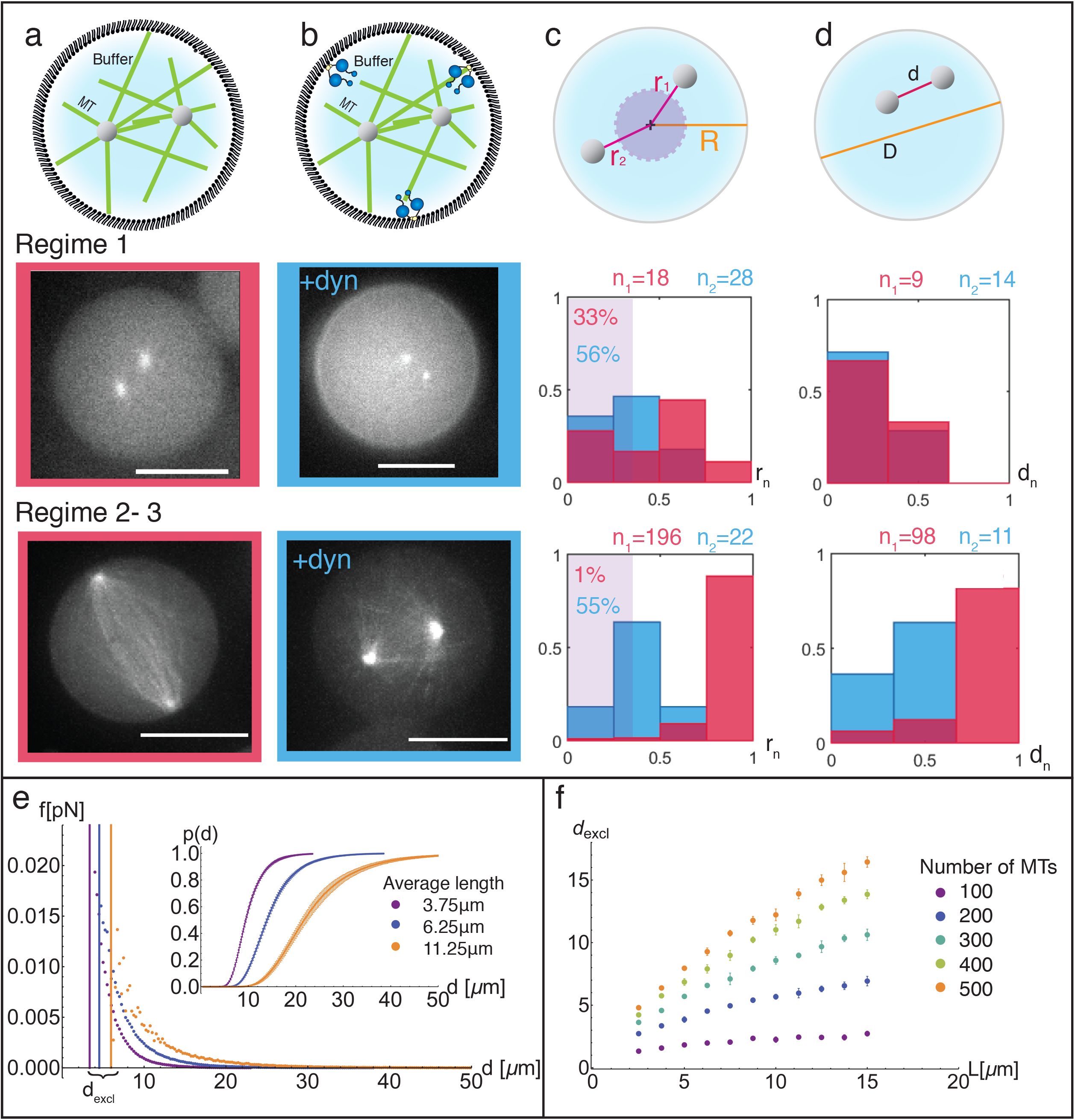
Double aster positioning: steric repulsive forces vs cortical centering forces. **(a)(b)** Positioning of two dynamic asters in the absence (a) and presence (b) of dynein at the cortex. Top: Schematic picture. Bottom: Spinning disk confocal fluorescence images for regimes 1 and 2-3, where the average MT length is respectively short (1) or intermediate/long (2-3) compared to the radius of the droplet. Scale=10μm. **(c)(d)** Quantification of aster position. Top: Representation of r, distance centrosome-centre, R radius of the droplet, d aster-aster separation, and D diameter of the droplet. Bottom: Quantification of 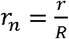 and 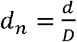 for asters in the absence (pink) or presence D (blue) of dynein. Bin size and normalization are as described in **Fig. 2**. For short MTs, asters are distributed throughout the sphere. For long MTs and no dynein, the two asters position at opposite poles. When dynein is present at the cortex, the asters position at moderate distance from each other. **(e)** Simulation results for steric repulsive forces (not considering confinement). The repulsive force *f* as a function of aster-aster separation d is plotted for different MT average lengths. The boundary of the hard core domain d< *d*_*excl*_, where asters cannot interpenetrate, is represented. Inset: Probability of insertion p(d) as a function of aster-aster separation for different MT average lengths. The number of MTs in each aster is 200. **(f)** Hard core domain size *d*_*excl*_ as a function of MT average length *L* for different MT numbers.

For a single aster, we determined the distance *r* to the centre of the droplet and normalized it to the droplet radius *R*: 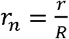. We defined an aster as centred if its centre was localized in a sphere of radius *r*_*n*_ = 0.36 concentric with the droplet, which contains 5% of the total droplet volume (**Fig. 2c**). In the absence of dynein at the cortex, when only pushing forces are present, we observed 15% of the centrosomes centred in regime 1 (N=73), 9% in regime 2 (N=74) and none in regime 3 (N=131) (**Fig. 2a and c**). Within spherical confinement, asters could thus be found centred only when the average MT length was shorter than the radius of the sphere, whereas they adopted a peripheral position when the average MT length increased, in accordance with previously published predictions^31^. We next introduced cortical pulling forces by anchoring dynein to the lipid monolayer^32, 33^. In the presence of pulling forces, we found most asters in a central position. Asters were found centred in 84% (N=93), 94% (N=18) and 71% (N=49) of cases for regimes 1, 2 and 3 respectively (**Fig. 2b and c**). We repeated these experiments with different batches of dynein and different lipid compositions and found similar results (**Supplementary Fig. S1bc**). Note however that stable centring is strictly dependent on the maintenance of end-on contacts between dynein and MT ends. As soon as sideways contacts are made, strong forces on individual MTs bring the aster to the periphery. This occurs most prominently at high concentrations of dynein and/or for long experimental times (**Supplementary Fig. S1c**). Interestingly, when asters were centred, the dynein distribution at the cortex was found to be homogeneous, whereas for peripheral asters clear colocalizing of dynein along MTs, consistent with sideways contacts, was observed (**Supplementary Fig. S1d**).

Next, we studied the positioning of two MT asters confined inside the same droplet. We determined both *r*_*n*_ and the normalized aster separation 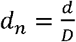 (*d* is the centre-to-centre aster separation and *D* the diameter of the droplet) for regimes 1 and 2-3 together. We found that in the absence of dynein, the presence of a second MT aster in the droplet did not drastically change the dependence of *r*_*n*_ on MT length (**Fig. 3a and 3c**). 33% of asters were found centred in regime 1 (N=18) compared to only 1% in regime 2-3 (N=196). We furthermore found in regime 2-3 that 67% of the aster pairs localized close to opposite sides of the droplet (*d*_*n*_ >0.8) (**Fig. 3a and 3d**, Supplementary movie 1). This suggests a repulsive force acting between asters. To elucidate the nature of the steric repulsion between two MT asters, we performed Monte Carlo simulations (Material and Methods-Simulations). We found that the repulsive potential and hence the force as a function of aster separation depends on the MT density and average length (**Fig. 3e**), but in all cases follows the same trend. For small inter-aster distances, we find a “hard-core” domain where the MTs coming from the other aster cannot penetrate. We defined *d*_*excl*_ as the minimum separation distance between the two asters (i.e. the diameter of this hard-core domain). *d*_*excl*_ increases with MT density for a given average MT length and also grows as a function of average length for a given density (**Fig. 3f**). For example, for 400 MTs, *d*_*excl*_ increases from 4μm for an average length of 3μm to 17μm for an average length of 15μm. Because the MTs splay out from the centrosome, leaving increasingly more free volume between them, it becomes easier to insert another aster at larger separations, which strongly decreases the repulsive force. In fact, we found the typical repulsion forces outside the hard-core domain to be ≲ 0.01pN, i.e. a hundred times smaller than typical forces generated by molecular motors or individual MTs.

We further investigated the positioning of two asters in the presence of cortical dynein and plotted *r*_*n*_ and *d*_*n*_ for regimes 1 and 2-3. We found 56% (N=28) of the asters centred in regime 1 and 55% (N=22) in regime 2-3 (**Fig. 3bc**). Similar to the single aster case, the presence of cortical dynein thus stabilized the central position. Intuitively, we had expected the presence of a second aster to prevent MTs from reaching the distal side of the droplet. In that case, the larger number of MTs captured by dynein from the proximal side would have pulled the aster to a peripheral position. Using the simulation data we therefore evaluated the probability *p*_*c*_ for MTs of one aster to cross over through the MTs of a second aster, as a function of aster separation *d* and angle θ (**Supplementary Fig. S2d**). For an aster separation distance of 4μm and for angles > 22.5^°^, the probability to cross over is already larger than 50% (computed for two asters containing 200 MTs each with a mean length of 6.25μm). MT crossovers are also experimentally visible (**Fig. 3b** and Supplementary movie 2). In regime 2-3, the proportion of droplets with an aster separation *d*_*n*_> 0.8 decreased from 67% without cortical dynein to 0% with cortical dynein (**Fig. 3d**). Apparently, the repulsive force between asters can be overcome by the presence of cortical dynein pulling on cross-over MTs. However, the most probable position was no longer in the middle of the droplet (**Fig. 3c**). We rather found the asters at a mean separation of *d*_*n*_= 0.34 (*sd* = 0.11), in line with the presence of a hard-core domain (d> *d*_*excl*_) as predicted by the simulations. This result shows that a basic spindle organization where two asters position at a “metaphase-like” distance from each other can readily be achieved by combining cortex-mediated centring forces with simple steric repulsion between asters.

To assess the effect of anti-parallel MT crosslinkers and motor proteins on the positioning of two asters, we next introduced the diffusive antiparallel cross-linker Ase1^35^, first alone and then in combination with kinesin 5 (both in the absence of dynein) (**Fig. 4a and 4c**). Again, we determined the aster position *r*_*n*_ (Supplementary Fig. S3) and the aster separation *d*_*n*_(**Fig. 4b**) for the different regimes in the presence of 0, 20, 36 and 80nM Ase1. For all Ase1 concentrations, the aster position *r*_*n*_ followed the same trend as before as a function of MT length. Asters started mainly centred in regime 1 and became more and more peripheral when MTs became longer. However, in regime 2-3 5 aster separation on average decreased when the concentration of Ase1 was increased (**Fig. 4b**). The population close to opposite poles (*d*_*n*_ > 0.8) decreased from 55% to 20%, 11% and 8% for 0, 20, 36 and 80nM Ase1 respectively. Accordingly, the population with an aster separation *d*_*n*_< 0.4 increased from 18% to 54%, 58% and 77%. The presence of Ase1 thus drastically reduces aster separation.

**Figure 4.**
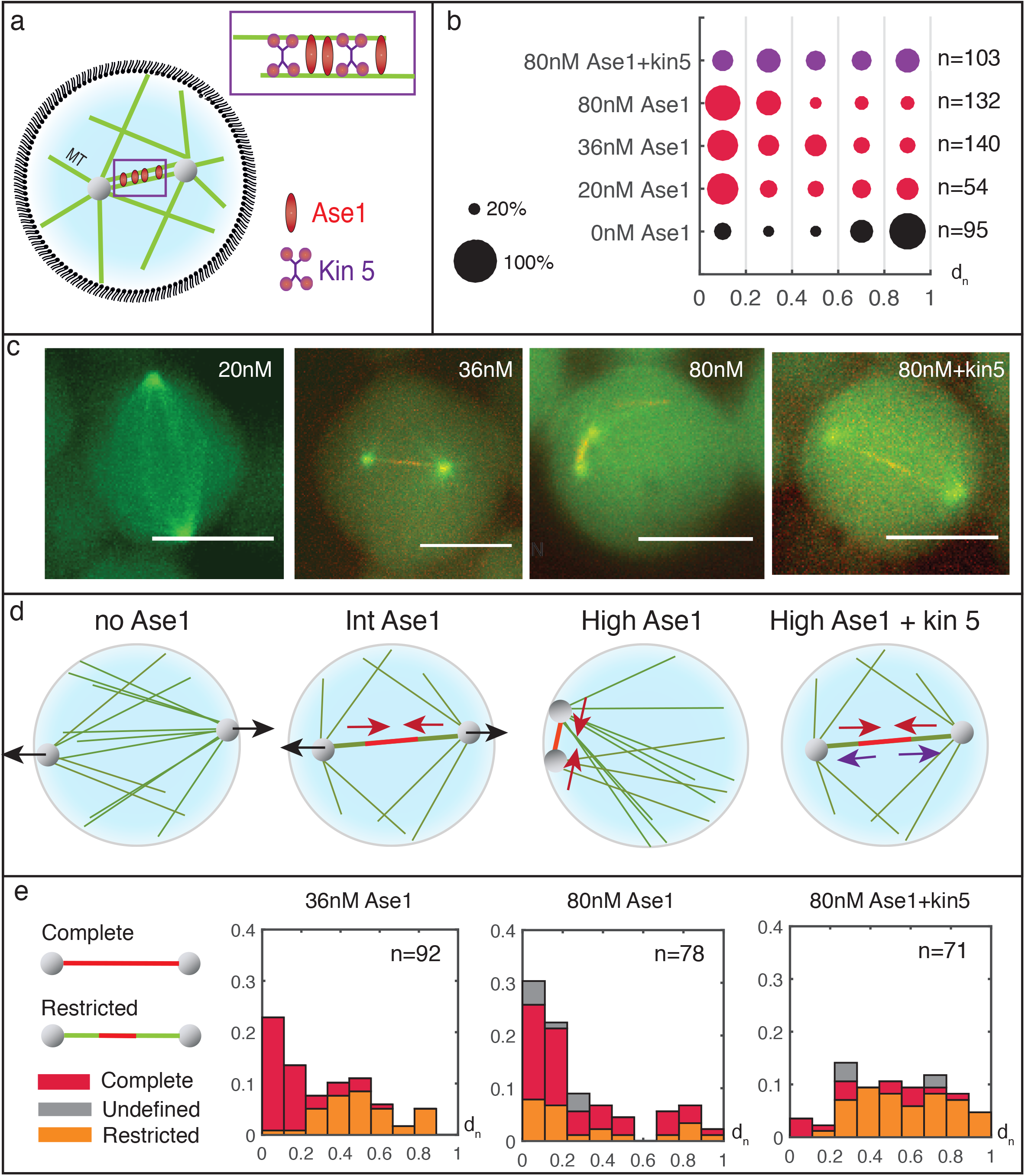
Aster-aster attraction by antiparallel crosslinks vs motor-driven repulsion. **(a)** Schematic picture of experimental components. **(b)** Percentile occurrence of aster-aster separation d_n_for droplets with intermediate or long MTs (Regime 2-3) for different Ase1 concentrations 0nM, 20nM, 36nM, 80nM, and 80nM Ase1 with Kinesin 5. **(c)** Spinning disk fluorescence confocal projections for different Ase1/kinesin 5 concentrations (MTs in green, Ase1/Kinesin5 in red, Scale=10μm). For 36nM and 80nM Ase1, and 80nM Ase1 with kinesin 5, the two centrosomes are bridged by a MT bundle enriched in Ase1/Kinesin5. For 80nM Ase1, the Ase1 signal reaches the two centrosomes, a situation referred to as a ‘complete’ link. For 36nM Ase1 and 80nM Ase1 with kinesin 5, the Ase1 signal is restricted to an overlap zone, referred to as a ‘restricted’ link. **(d)** Cartoon of the forces involved in centrosome separation for the cases: no Ase1 (0nM Ase1), Int Ase1 (36nM Ase1), High Ase1 (80nM Ase1) and High Ase1 + kin5 (80nM Ase1 + kinesin 5). Black, red and purple arrows represent respectively steric repulsive, Ase1-driven attractive and motor-driven repulsive forces. **(e)** Quantification of Ase1 links for a subset of data presented in (b). Only data presenting a clear Ase1 enriched bundle bridging the two centrosomes are included. The fraction of droplets with a complete or restricted link and with d_n_in a given range is plotted.

Often, we noticed the presence of Ase1-enriched MT bundles bridging the centrosomes (**Fig. 4c**). Such bundles were clearly seen in 13%, 63% and 58% of the cases for 20, 36 and 80nM Ase1 respectively. We distinguished two types of Ase1-bundles: complete and restricted bundles (**Fig. 4e**). Complete bundles showed an Ase1 signal reaching from one aster all the way to the other while in restricted bundles, Ase1 localized to a restricted section in the middle of the bundle. For 36 and 80 nM Ase1, in cases where bundles were clearly seen, we again plotted *d*_*n*_, indicating this time also the nature of the bundle (**Fig 4e**). For 80 nM Ase1, we found 60% of the population at small aster separation (*d*_*n*_ < 0.2), with a majority of complete bundles. For 36 nM, the population found at aster separation *d*_*n*_< 0.2 decreased to 47% and a second population at higher aster separation *d*_*n*_ showing mainly restricted bundles appeared. Apparently, at higher concentrations, Ase1 is increasingly able to overcome the aster-aster repulsion forces leading to a reduction of aster separation. This could either be the result of stabilization of anti-parallel MT bundles early on in the process of aster positioning (when MTs are still short and centrosomes not yet separated), or due to entropic forces leading to inward sliding between anti-parallel MTs^30^. Our experiments monitor the steady state outcome of this process and can therefore not distinguish between these two possibilities.

We finally added kinesin 5 in the presence of 80nM Ase1. Here also, asters gradually adopted a peripheral position with increasing average MT length (Supplementary Fig. S3). However, in regime 2-3 the presence of kinesin 5 increased the aster separation *d*_*n*_again, compared to the situation with Ase1 alone (**Fig. 4b**). In cases where Ase1-rich bundles could clearly be seen, we now predominantly observed restricted bundles (**Fig. 4e**). Restricted bundles thus seem the result of a competition between Ase1-mediated inward forces and kinesin 5-mediated outward forces. The presence of restricted bundles at moderate Ase1 concentrations in experiments without kinesin 5 suggests that Ase1-mediated inward forces can alternatively be balanced by outward forces due to aster-aster repulsion.

In this letter we show that two dynamic MT asters are able to repel each other without any additional MAPs, leading to spindle pole separation. It was earlier proposed that MTs pushing directly against the opposite centrosome may drive centrosome separation during prophase^36^. We show however that asters exhibit a strong repulsive core that depends on the density and mean length of MTs, and is mainly due to MT-MT interactions rather than direct MT-centrosome contacts (**Supplementary Fig. S2c**). *In vivo*, the number of nucleated MTs per centrosome varies drastically with cell type. For a mean MT length of 10μm, 500 MTs would be sufficient for *d*_*excl*_ to reach 12μm according to our simulations. *In vivo*, this repulsion force might even be stronger because of several reasons. First of all, MTs are subject to thermal fluctuations, effectively occupying a larger volume than their rod-shaped molecular volume. MTs are also nucleated within the spindle^38^, increasing the local density of MTs further away from the centrosome, thus increasing the local repulsion force. It is further interesting to note that during mitosis the number of centrosome nucleated MTs can increase up to 5-fold^37^. The control of both local and global MT density may thus contribute to bipolar spindle organization.

We further show that the repulsive force between asters can alternatively be balanced by cortical dynein-mediated centring forces or by Ase1-mediated attractive forces in MT overlaps. In cells, cortical dynein is often implicated in pulling the spindle poles apart^5, 39–41^. For this to work, the MT distribution needs to be such that MTs are preferentially captured at the proximal side of the aster. It was earlier suggested that the physical presence of two centrosomes close together might be enough to prevent MTs from growing towards the other side^42^. Our results suggest that additional mechanisms much be in place, such as depolymerisation of MTs or additional barriers for MT growth at the midzone, or regulation of the spatial distribution of cortical dynein.

Previously, aster separation has been thought to arise from the activity of molecular motors on antiparallel MTs in the spindle midzone. One well-known model is the “push-pull” model^43^, where centrosome separation results from the antagonistic activity of motors in the overlap. Our results show that the presence of the passive diffusible cross-linker Ase1 can also contribute to aster separation. Lansky et al^30^ previously showed that Ase1 can generate pN entropic forces *in vitro*, which is sufficient to antagonize kinesin14 Ncd activity in MT overlaps. These entropic forces lead to an attractive force between centrosomes in our experiments. We found that kinesin 5 motors were able to overcome the action of Ase1. Kinesin 5 generates 5-7 pN forces per motor^44^, notably stronger than kinesin 14, which can generate about 0,1 pN per motor^45^.

In conclusion, we have presented the *in vitro* reconstitution of minimal spindles as a system where the contribution of spindle components to overall spindle organization can be assessed in a direct way (**Fig. 5**). We find that MT asters inherently repel each other as MT length increases, leading to a default “anaphase-like” aster positioning in confined spaces. This repulsive force can be counteracted to produce “metaphase-like” aster positioning using different molecular strategies, which in turn can be counteracted to recover anaphase-like positioning by adding additional components. An interesting task for the future will be to challenge the robustness of these different molecular strategies by the addition of other essential spindle components (such as artificial chromosomes) and to test what level of molecular complexity would be required for controlled transitions between different spindle configurations, as would be relevant for the progress of mitotic spindles through the cell cycle in real cells.

**Figure 5.**
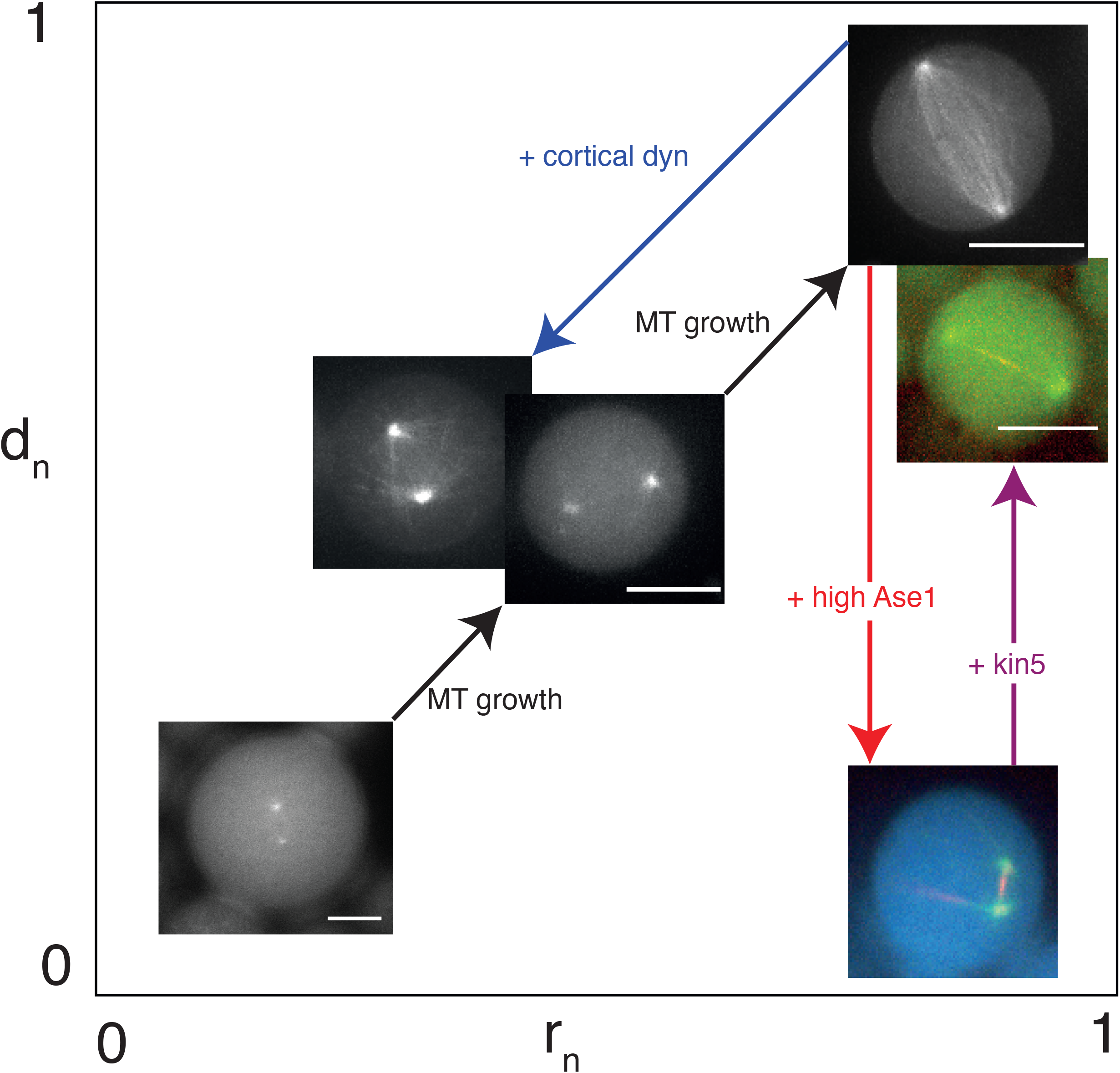
Schematic diagram of spindle configurations. Images of typical spindle configurations as observed in our experiments are represented in an aster position (r_n_) versus separation (d_n_) “plane”. The effects of increasing MT length (black arrows) and/or addition of cortical dynein, Ase 1 and Kin5 (coloured arrows) on spindle configuration are indicated.

## Acknowledgements

We thank: Samantha Reck-Peterson’s group and in particular Sirui Zou for help with dynein purification, Patrick Tabeling’s group and in particular Bingqing Shen for initial help with microfluidics, Marcel Janson for providing Ase1 protien, and Zdenek Lansky for providing kinesin5 protein. The work of SR, MV and MD was supported by funding from the European Research Council (ERC Synergy grant MODEL CELL). ICG was financially supported by the FOM (Foundation for Fundamental Research on Matter) programme 110 “Spatial design of biochemical regulation networks”. The work of BMM is part of the research programme of the Netherlands Organisation for Scientific Research (NWO) and was performed at the research institute AMOLF.

