## Supplementary material for "Reconstitution of basic mitotic spindles in cell-like confinement": Methods

### Materials and methods

### Experiments Reagents and proteins

### Lipids and surfactants: 1,2-dioleoyl-*sn-*glycero-3-phospho-l-serine (sodium salt) (DOPS), 1,2-dipalmitoyl-*sn*-glycero-3-phosphoethanolamine-N-(biotinyl) (biotin PE) were purchased from Avanti Polar Lipids, (Alabaster, AL). Span^®^80 was purchased from Sigma Aldrich (USA). Buffers: Reagents for MT assays were dissolved in MRB80 buffer (80mM K-pipes, 4mM MgCl2, 1mM EGTA, pH 6.8) and were obtained from Sigma, unless otherwise stated. Tubulin was purchased from Cytoskeleton Inc (Denver, USA).

### Purified proteins: The dynein construct used was the biotinylated GST-dynein331 (referred to as dynein) which contains a GFP, a SNAP tag and a Halotag. Details of the construct and the purification procedure are described elsewhere^1^^,^ ^2^. Purified GST-dynein-331 was biotinylated on the SNAP tag (SNAP biotin®, New England Biolabs) and labelled with a fluorophore via its Halotag (Promega). Three batches of dynein were used for this paper. Batch 1 was used for single and double aster positioning presented in Fig. 2 and Fig. 3. Batch 2 was used for single and double aster positioning presented in Supplementary Fig. S2b. Batch 3 was used for the experiments presented in Supplementary Fig. S1c and the pictures presented in Supplementary Fig. S1d.

### Centrosomes were purified from human lymphoblastic KL37 cell lines (Leibniz-Institut DSMZ, ACC 46) following a published protocol ^3^. Recombinant histidine-tagged full length S.pombe GFP-Ase1 (referred to as Ase1) was kindly provided by Janson lab (Cell Biology, Wageningen University, the Netherlands). Recombinant GFP-Cut7, S.pombe Kinesin5 (referred to as kinesin5), was kindly provided by Zdenek Lansky, Diez lab (Max-Planck Institute of Molecular Cell Biology and Genetics, Dresden, Germany).

Oil-lipid mixture

Lipids (DOPS, biotin PE) were mixed in chloroform, dried under nitrogen flow, placed under vacuum for 30min and dissolved in mineral oil (M5904, Sigma-Aldrich). Span^®^80 was subsequently added to the mixture and sonicated for 30 min. Two oil/lipid/surfactant mixture were used in this paper. Mix 1 contained lipids at a molar ratio DOPS/biotin PE 99:1 with a concentration 0,7mg/mL and Span^®^80 at 3,1wt% in mineral oil. Mix 2 contained lipids at a molar ratio DOPS/biotin PE 58:41 with a concentration 1,4mg/mL and Span^®^80 at 3,1wt% in mineral oil. Mix 1 was used for single and double aster positioning in presence/absence of cortical dynein in Figure 2 and 3. Mix 2 was used for and all subsequent experiments including Supplementary Fig. S1 and S2. We confirmed the cortical position of dynein for both conditions, as well as the bipolar organization of two dynamic asters in confinement (**Supplementary Fig. S1 and S2**).

MT mix

MT mix for experiments with oil-lipid mix 1 : Tubulin (25-30 uM), Fluorescent tubulin (Hilyte 488) from porcine brain (2.5-3 uM), GTP (5 mM), Streptavidin (40nM), ATP (1 mM), Fluorescent Dextran (10,000MW, Alexa Fluor® 647) (20μM), purified centrosomes (concentration adjusted to have mainly one or two centrosomes per droplet), Glucose (50mM) with an Oxygen Scavenging system (Glucose oxidase 0.1 mg/mL, DTT 4 mM, catalase 0.1 mg/mL) and an ATP regenerating system (Phospho(enol)pyruvic acid monosodium salt hydrate (PEP) 26.6 mM, Pyruvate kinase/lactic dehydrogenase (PK/LDH) 26.6units) supplemented optionally with dynein (TMR labeled, X3).

MT mix for experiments with oil-lipid mix 2 : same as above except Fluorescent tubulin (Rhodamine), Streptavidin (200nM), Fluorescent Dextran (0.6μm) and addition of Bovine serum albumin (2mg/mL), Tween (0.06%). Mix was supplemented optionally with dynein, Ase1 (20,36,80nM) and/or kinesin 5 for midzone experiments.

The mix was centrifuged (Airfuge® Air-driven ultracentrifuge, Beckman Coulter, USA) at 4◦C for 5 min at 30 psi (200,000 rcf ) without the centrosomes. Centrosomes were then added and mixed gently.
For kinetics comparisons (Supplementary Fig. S1c, right) with/without cortical dynein, a common MT mix was prepared, centrifuged, and centrosome added. The mix was then separated in two. One part was supplemented with dynein, the other with the same amount of MRB80.

Emulsion droplets formation and Imaging

Emulsion droplets of MT mix in oil/lipid mixture were prepared using a flow-focusing microfluidic chip as described in ^1^^,^ ^4^. Liquids were driven with a pressure controller MFCS-FLEX-4C-1,000 mbar (Fluigent, Paris) and PEEK tubing (Cluzeau Info. Labo, France). Droplet formation was observed with a Leica PMIRB-inverted microscope in bright field and flow rate were adjusted to obtain droplets around 15-25 µm diameter (see histogram below). Droplets were then collected in a flow cell coated with Momentive RTV615 (Lubribond, The Netherlands). Droplets presenting one or two centrosomes were imaged with a spinning disk confocal head CSU-X1 from Yokogawa (Japan) and a cooled EM-CCD camera iXon3 (Andor, UK) at 26⁰C with a 100X oil immersion objective and excitation lasers 488, 561 and 641 nm (Andor, UK). Stacks were taken with 1µm intervals (~20 images/droplets). For kinetics comparisons with/without cortical dynein (**Supplementary Fig. S1c, right**), two droplet sets coming from the same MT mix as described in the paragraph above were formed and kept on ice. The two emulsions were then introduced in two adjacent flow channels for imaging. Images were taken alternating between both channels, revealing the evolution overtime of the two samples.

|  | % |
| --- | --- |
| 10<D<15 | 1.28996 |
| 15<D<20 | 51.5984 |
| 20<D<25 | 34.0437 |
| 25<D<30 | 8.13236 |
| 30<D<35 | 1.9069 |
| 35<D<40 | 0.616938 |

Figure 1: Histogram of Droplet diameter (µm) for all the experiment shown in this paper.

Image analysis:

Droplet 3D shape was determined from the fluorescent dextran z-stack thanks to a homemade matlab program. Due to optical effects coming from imaging an aqueous droplet in oil, only z slices around the equator were taken into account, binarized, and an ellipse 2D fitted. To reconstruct the droplet 3D shape, an ellipsoid 3D was then fitted using the least square optimization technique adapted from the program ‘Ellipsoid fit’ developed by Yuri Petrov. Adequation of the 3D fit with the z-stack images was verified. The centre coordinates and mean radius “R” of the droplet were recorded. Thanks to a homemade matlab GUI, the centrosome position was determined in the fluorescent tubulin z-stack. From the 3D coordinates of the centrosomes and the centre of the droplet, the distance to the center $r$ was calculated and normalized $r_{n}=\frac{r}{R}$. In the case where two centrosomes were present in one droplet, the separation $d$ between the two centrosomes was also determined and normalized, $d_{n}=\frac{d}{D}$.

**Simulations:**
Aster modelling:

Since the persistence length of a microtubule ^5^ is two orders of magnitude larger than the typical diameter of the emulsion droplets, we can model them as rigid rods, and chose to represent them as spherocylinders with a diameter of $D=25 nm$. The centrosome is represented by a hard sphere with radius $R_{c}=0.25 \mu m$, from which $N$ MTs project in the radial direction. The orientation of the MTs is chosen from an isotropic distribution and their lengths are exponentially distributed. We have considered asters containing between $N=100$ and $N=600$MTs, the upper bound being set by the total amount of tubulin available in a typical-sized emulsion droplet. The mean lengths considered varied between $L_{m}=2.5 \mu m$ and $L_{m}=15 \mu m$.

Interaction geometry:

The repulsion force between the two asters, being of steric origin, only acts within the lens-shaped volume where actual physical contact between the two asters can occur. This lens-shaped volume is defined as the intersection of the bounding spheres surrounding the asters, each bounding sphere being centred on the respective centrosome and having a radius$R_{b}=L_{max}+R_{c}$, where $L_{max}$ is the length of the longest MT in the aster. The volume is situated symmetrically in the midpoint between the two centrosomes and, except in extreme cases, is not intersected by the confining surface. This allows us to decouple the aster-aster repulsion from the cortex related positioning. We therefore study the repulsion between two asters in bulk, without the presence of the confining cortex.

Repulsive force calculation:

The asters are too complex geometrical objects to allow an analytical calculation of their excluded volume. However, similar to what was done earlier for polymer coils^6^, it is possible to estimate the effective repulsion potential between two such objects from the so-called insertion probability $p\left( d \right),$ being the probability that one can insert an object at a distance $d$ from the center of mass of the second one without creating an overlap between the objects. Statistical mechanical arguments^7^ show that this probability is related to potential of mean force between the two objects, through the relation $U\left( d \right)=-k_{b}Tln(p(d))$, where $k_{b}$ is Boltzmann’s constant and $T$ the absolute temperature, here taken to be room temperature.
