## Supplementary Fig and Movie Legends for "Reconstitution of basic mitotic spindles in cell-like confinement"

**Supplementary Figure legends**

**Figure S1:** Single aster positioning
(a) Dynein localization to the cortex for Oil-lipid mixture Mix 1 (left) and Mix2 (right). Mix1: Dynein localization at the cortex is verified by the addition of short MT seeds in the mix. In the case of “no dynein”, MT seeds are localized inside the droplet. When dynein is present, MT seeds are captured at the cortex. Mix 2: Dynein localization at the cortex is verified by fluorescent labelling. Left picture shows fluorescent streptavidin. Right picture shows fluorescent dynein (red), when the aster (white) is centered.
(b) Positioning of one dynamic aster in the absence (pink) or presence (blue) of dynein at the cortex with dynein from dynein batch2 (18nM) and oil-lipid mixture Mix2.
(c) Left: Percentage of centred centrosomes for MTs in regime 2 (L2) and 3 (L3) at different dynein concentrations 0nM, 10nM, 20nM, 30nM, 60nM (batch3, 1 experiment/ concentration). Right: Kinetic comparison of aster positioning with (blue)/without (pink) dynein (dynein batch 3, 30nM) at the cortex shows that dynein presence at the cortex of the droplet delays the ultimate peripheral position of the aster.
(d) Spinning disk confocal fluorescence image of dynein at the cortex. Left: dynein is homogeneously distributed at the cortex. Aster is centred. Middle: Dynein starts to colocalize with MT at the cortex (red arrow). Right: Dynein is colocalizing with MTs at the cortex. Aster is peripheral.
 **Figure S2:** Double aster positioning
(a) Positioning of two dynamic asters in absence (pink) or presence (blue) of dynein at the cortex with dynein from batch2 (18nM) and oil-lipid mixture Mix2.
(b) Schematic representation of the forces acting during double aster positioning with/without dynein. Top: without dynein, asters repulsive force (black arrows) leads them as far as possible in the confinement. Bottom: with dynein at the cortex, asters repulsive force (black arrows) compete with dynein pulling force (blue arrows). The blue arrows are centering only if more MTs are pulling from the distal side than the proximal side.
(c) Simulation: Number of MT-MT and MT-centrosome overlaps as a function of aster separation shows that the repulsive force is mainly due to MT-MT steric interaction rather than MT-centrosome. Each aster contains 200 MTs with a mean length of 6.25µm.
(d) Simulation: Probability $p_{c}$ for a MT with angle θ to cross over as a function of aster separation $d$.

Parameters used are the same as for panel as for (c).

**Figure S3:** Double aster positioning in presence of Ase1 and kinesin5

Percentage of asters centred as a function of MT regime and for different concentrations of Ase1 with/without kinesin5. All the curves follow the same trend. Asters adopt a peripheral position as MT grow.

**Supplementary Movie legends**

**Movie S1:** MT cytoskeleton self-organize in a bipolar structure when confined in a 3D sphere (3D reconstruction of spinning disk confocal fluorescence microscopy).

**Movie S2:** Double aster confined in 3D w/o droplet show MTs crossing over and reaching the distal side of the droplet, even in the presence of the second aster (3D reconstruction of spinning disk confocal fluorescence microscopy).
