## Supplementary figures and images for "Reconstitution of basic mitotic spindles in cell-like confinement"

### Supp Figure 1

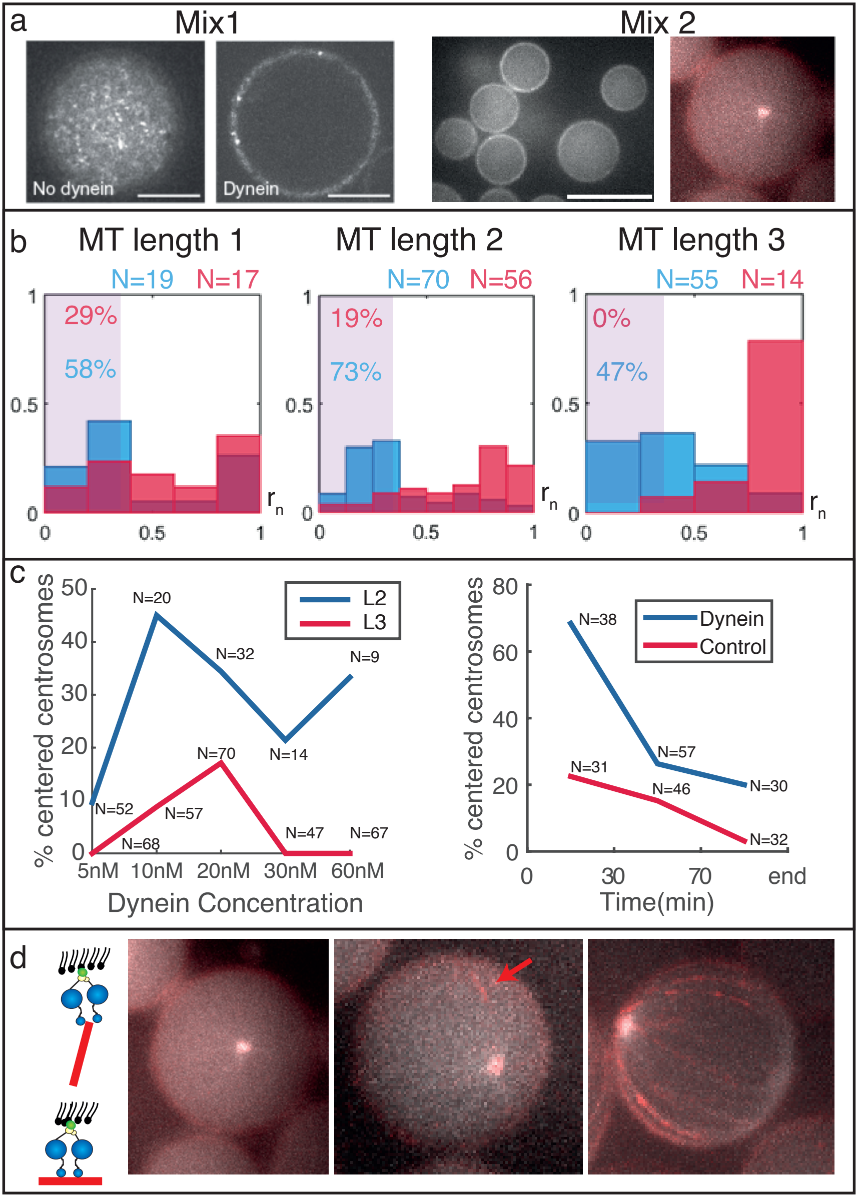

### Supp Figure 2

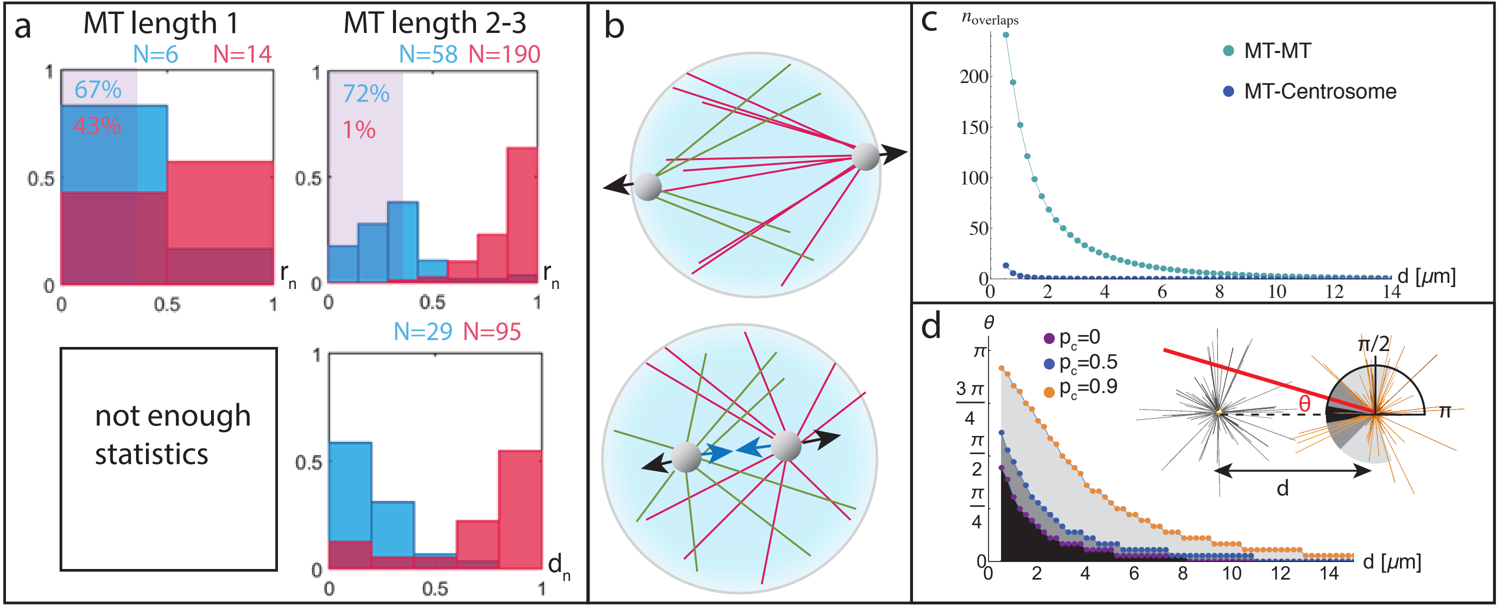

### Supp Figure 3

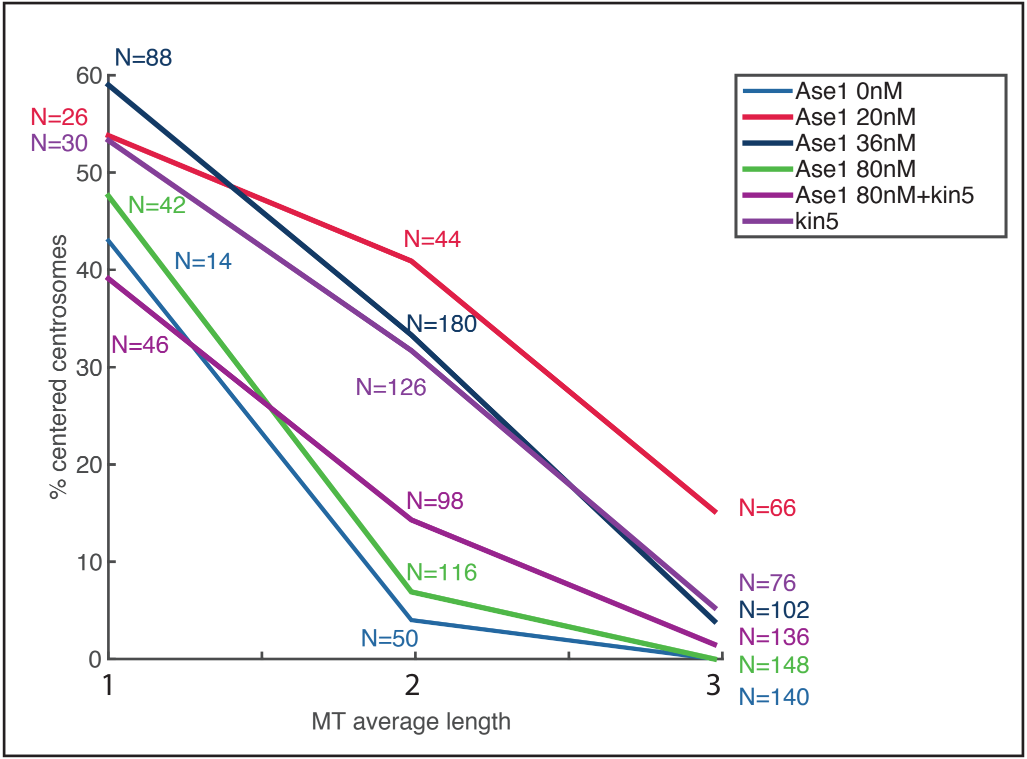
